# The roles of LINEs, LTRs and SINEs in lineage-specific gene family expansions in the human and mouse genomes

**DOI:** 10.1101/042309

**Authors:** Václav Janoušek, Christina M. Laukaitis, Alexey Yanchukov, Robert C. Karn

**Affiliations:** Department of Zoology, Faculty of Science, Charles University in Prague, Czech Republic.; Institute of Vertebrate Biology, ASCR, Brno, Czech Republic.; Department of Medicine, College of Medicine, University of Arizona, Tucson, AZ, USA.; Department of Biology, Faculty of Arts and Sciences, Bülent Ecevit Üniversity, Zonguldak, Turkey.

**Keywords:** Gene families, transposable elements, retrotransposons, LINE, LTR, SINE, human, mouse

## Abstract

We explored genome-wide patterns of RT content surrounding lineage-specific gene family expansions in the human and mouse genomes. Our results suggest that the size of a gene family is an important predictor of the RT distribution in close proximity to the family members. The distribution differs considerably between the three most common RT classes (LINEs, LTRs and SINEs). LINEs and LTRs tend to be more abundant around genes of multi-copy gene families, whereas SINEs tend to be depleted around such genes. Detailed analysis of the distribution and diversity of LINEs and LTRs with respect to gene family size suggests that each has a distinct involvement in gene family expansion. LTRs are associated with open chromatin sites surrounding the gene families, supporting their involvement in gene regulation, whereas LINEs may play a structural role, promoting gene duplication. This suggests that gene family expansions, especially in the mouse genome, might undergo two phases, the first is characterized by elevated deposition of LTRs and their utilization in reshaping gene regulatory networks. The second phase is characterized by rapid gene family expansion due to continuous accumulation of LINEs and it appears that, in some instances at least, this could become a runaway process. We provide an example in which this has happened and we present a simulation supporting the possibility of the runaway process. Our observations also suggest that specific differences exist in this gene family expansion process between human and mouse genomes.

## INTRODUCTION

One of the surprises that emerged from the draft sequence of the human genome was that half or more of it is composed of interspersed repetitive DNA sequences that were thought to be parasitic DNA in some state of degradation (Lander et al. 2001). These transposable elements (TEs) were first discovered by McClintock 70 years ago (see for example (McClintock 1950)). TEs are mobile DNA sequences that can move from one site in a genome to another. On arrival, they either insert themselves directly into the genomic DNA by a cut-and-paste mechanism (transposons) or indirectly through an RNA intermediate (retrotransposons; RTs). Since their discovery, the numbers and kinds of TEs that have been described have grown into a complex collection that warranted a classification system (Wicker et al. 2007; Kapitonov and Jurka 2008). Since the TEs in mammal genomes are mostly retrotransposons: LINEs, LTRs and SINEs, we will refer to these collectively as RTs.

The idea that TEs are nothing more than parasitic DNA that infiltrated eukaryotic genomes has been challenged recently with the suggestion that they have played an important role in genome evolution (reviewed in(Fedoroff 2012)). In fact, McClintock’s observation that TEs can control gene expression (McClintock 1950; McClintock 1956) presaged recognition of their evolutionary involvement in the architecture of gene regulatory networks (Feschotte 2008; Bourque 2009). Since then, TEs have been found to contain functional binding sites for transcription factors (Jordan et al. 2003; Bourque et al. 2008; Polavarapu et al. 2008; Sundaram et al. 2014) and recently, DNAse I hypersensitive site (DHS) data from ENCODE were used to show that 44% of open chromatin regions in the human genome are in RTs, as are 63% of regions controlling primate-specific gene expression (Jacques et al. 2013). TEs, particularly ERVs, have contributed hundreds of thousands of novel regulatory elements to the primate lineage and reshaped the human transcriptional landscape (Jacques et al. 2013). Genes proximal to tissue-specific hypomethylated RTs are enriched for functions performed in that tissue, (Xie et al. 2013), emphasizing the importance of RTs in contributing to regulating tissue-specificity of gene expression in the mouse. Using a ChIP-seq approach to map binding sites of 26 orthologous transcription factors (TFs) in the human and mouse genomes RTs have been found to contribute up to 40 % of some TF binding sites, most of which were species-specific with some binding sites being significantly expanded only in one lineage (Sundaram et al. 2014).

Besides their importance in gene regulation, RTs are also considered to be an important source of structural variation (Bourque 2009). RTs may provide homologous substrates for double-strand break (DSB) induced repair mechanisms, including non-allelic homologous recombination (NAHR) and microhomology-mediated break-induced repair (MMBIR), which may result in structural variation (Hastings et al. 2009). Double-strand breaks themselves may be associated with repetitive elements (Hedges and Deininger 2007; Argueso et al. 2008). Accordingly, segmental duplications and CNVs were repeatedly found to have TEs enriched at their edges (Bailey et al. 2003; Kim et al. 2008; She et al. 2008). Some studies confirmed directly the role of TEs in NAHR (Fitch et al. 1991; Janoušek et al. 2013; Campbell et al. 2014; Startek et al. 2015).

Now, a new view of the complex role of TEs in organismal evolution is being adopted, suggesting that TE mobilization may represent an important source of new genetic variability under stressful conditions (Capy et al. 2000; Fablet and Vieira 2011). A role for TEs has been proposed in adaptive evolution of an invasive species of ant, *Cadiocondyla obscurior* (Schrader et al. 2014). This species has a small genome with rapidly-evolving accumulations of TEs, called TE islands. The species produces genetically-depleted founder populations (reviewed in (Stapley et al. 2015)). When the genomes of two isolated populations of *C. obscurior* were compared, distinct phenotypic differences were found between them with a strong correlation between the TE islands and genetic variation, suggesting that these serve as a source of variation in the founder populations. The origin of repetitive elements often correlates with speciation events, suggesting that TEs might have played major roles in evolution, and possibly speciation (Jurka et al. 2011). (Fedoroff 2012) suggested that the evolution of a powerful epigenetic apparatus enabled a proliferation of TEs and their successful co-option in the evolution of the high complexity of eukaryote genomes.

Since Ohno’s proposal that gene duplication represents an important source of new genetic material (Ohno 1970), evidence for its importance in adaptation to changes in the environment has mounted (reviewed in (Kondrashov 2012)). Because gene duplication provides a means for gene family expansion and thus the production of new genetic material (Korbel et al. 2008), we feel that it is time to further explore the role of TEs in the evolution of gene families. In a previous study (Janoušek et al. 2013), we examined the role of repeat element sequences in the expansions of the mouse and rat *Abp* gene families and found high densities of L1 and ERVII repeats in the *Abp* gene region with abrupt transitions at the region boundaries, suggesting that their higher densities are tightly associated with *Abp* gene duplication. We confirmed the role of L1 repeats with the identification of recombinant L1Md_T elements at the edges of the most recent mouse *Abp* gene duplication, suggesting that they served as the substrates for NAHR. We observed that the major accumulation of L1 elements occurred after the split of the mouse and rat lineages and that there is a striking overlap between the timing of L1 accumulation and the expansion of the *Abp* gene family in the mouse genome. This established a link between the accumulation of L1 elements and the expansion of the *Abp* gene family, and identified an NAHR-related breakpoint in the most recent duplication. At that time, the reason for the large accumulation of ERVII elements that occurred before the gene family expansion was not obvious.

In the study we report here, we endeavored to determine how widespread is the involvement of RTs in human and mouse gene family expansions, and what putative roles these elements play in gene family evolution. We found a significant association between RT content and the size of lineage-specific gene family expansions, and LINEs and LTRs were found to have an important role in these. Detailed analysis revealed the complex role these elements play and we propose a model of interaction of LINEs and LTRs supported by the *Abp* gene family example. We also suggest that gene family expansions, especially in the mouse genome, apparently occur in two phases. The first phase is characterized by elevated deposition of LTRs and rewiring gene regulatory networks due to an increase in number of gene copies. The second phase is then characterized by continuing rapid expansion due to ongoing accumulation of LINEs, potentially becoming a runaway process. We constructed a computer simulation to investigate the theoretical mechanisms that could allow this second phase to assume runaway proportions.

## RESULTS

### Retrotransposon content and gene family size

We explored the densities, abundances and lengths of the three main classes of retrotransposons (RTs, which include LINEs, LTRs and SINEs) as a function of gene family size in the human and mouse genomes. The retrotransposons analyzed were active only after the mouse-human split and were thus lineage-specific. We began by assessing the importance of gene family size and window size in explaining RT densities, abundances, and average lengths in the two species separately. In these considerations, gene family sizes ranged from single genes to inparalog/outparalog numbers >10 genes. We tested window sizes of 10 Kb, 50 Kb, 100 Kb, 500 Kb, 1 Mb, and 5 Mb surrounding the inparalogs/outparalogs of individual gene families. The effect of gene family size and window size and their interaction (i.e. the full model) was found to explain the data best of all possible tested models for all three RT classes in both species for the inparalog and outparalog datasets (**Table 2; Supplemental Table 2**). Akaike’s information criterion (AIC; (Akaike 1974)) analysis exhibits the lowest values for the full model for all tested datasets. The estimated parameters of the full model were plotted in **Fig. 1A**.

**Figure 1.**
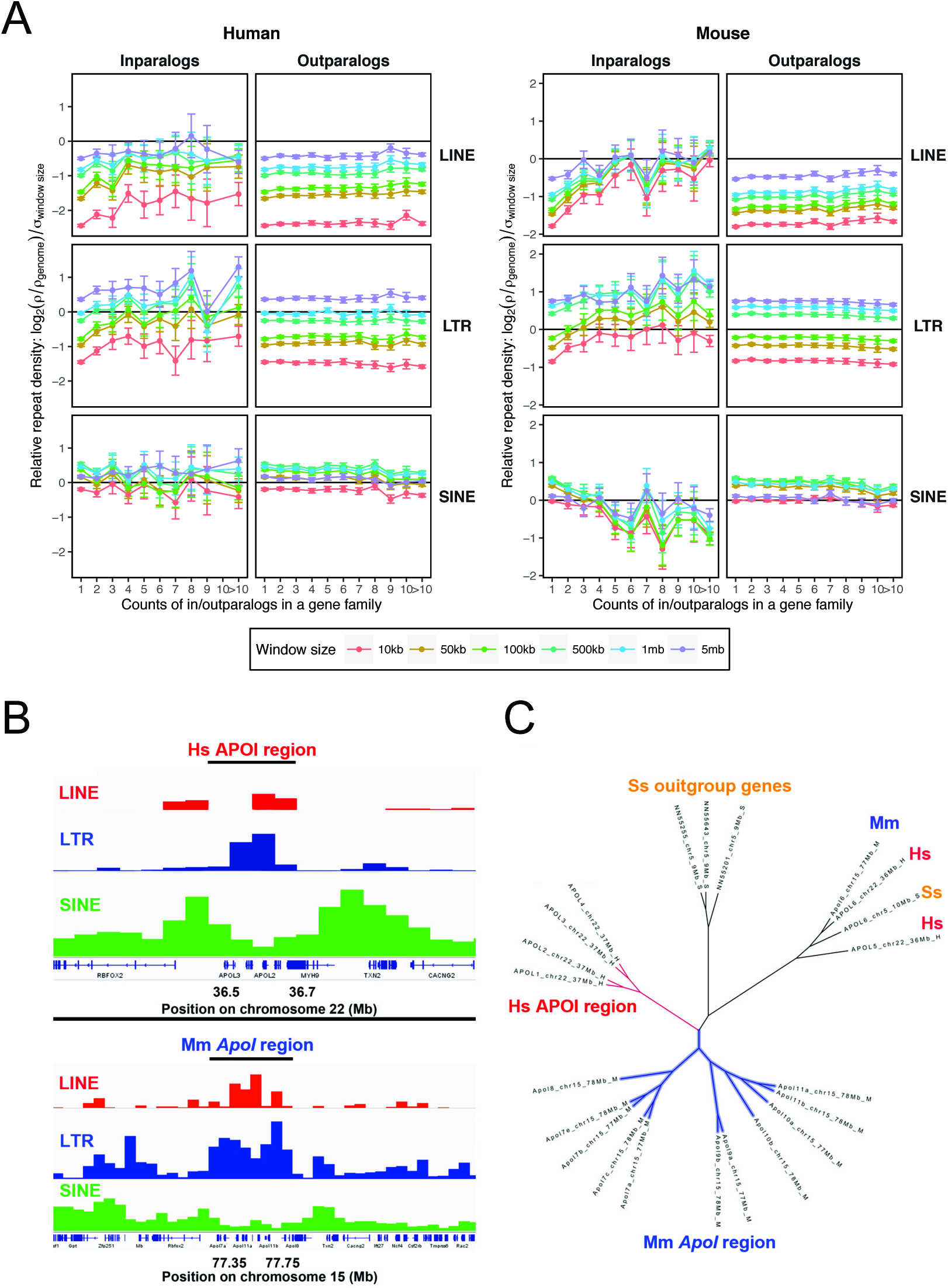
(A) The relative densities of the lineage-specific RT classes (LINE, LTR and SINE) around the genes from gene families of increasing size in the human (left) and the mouse (right) genomes. Gene family size is defined as the number of inparalogs/outparalogs in a gene family with size greater than one, while the gene family size of one corresponds to single genes. The RT densities represent the proportion of base pairs contributed by an RT class in a given window size (10 Kb, 50 Kb, 100 Kb, 500 Kb, 1 Mb, 5 Mb) around single genes, inparalogs and/or outparalogs of individual gene families. The densities are scaled so that the zero corresponds to the genome-wide average for an RT class and window size and data were treated for differing variances among different size windows. Positive and negative values represent RT densities higher and lower than the genome-wide average, respectively. Panels (B) and (C) show a representative gene family, *ApoI*, in the human and the mouse genomes. Panel B compares the gene family sizes and RT content in the two taxa and Panel C shows a gene tree of the human (red), mouse (blue) and pig (*Sus scrofa*) genes (brown), which was used to infer mouse and human lineage-specific duplications (inparalogs).

**Figure 2.**
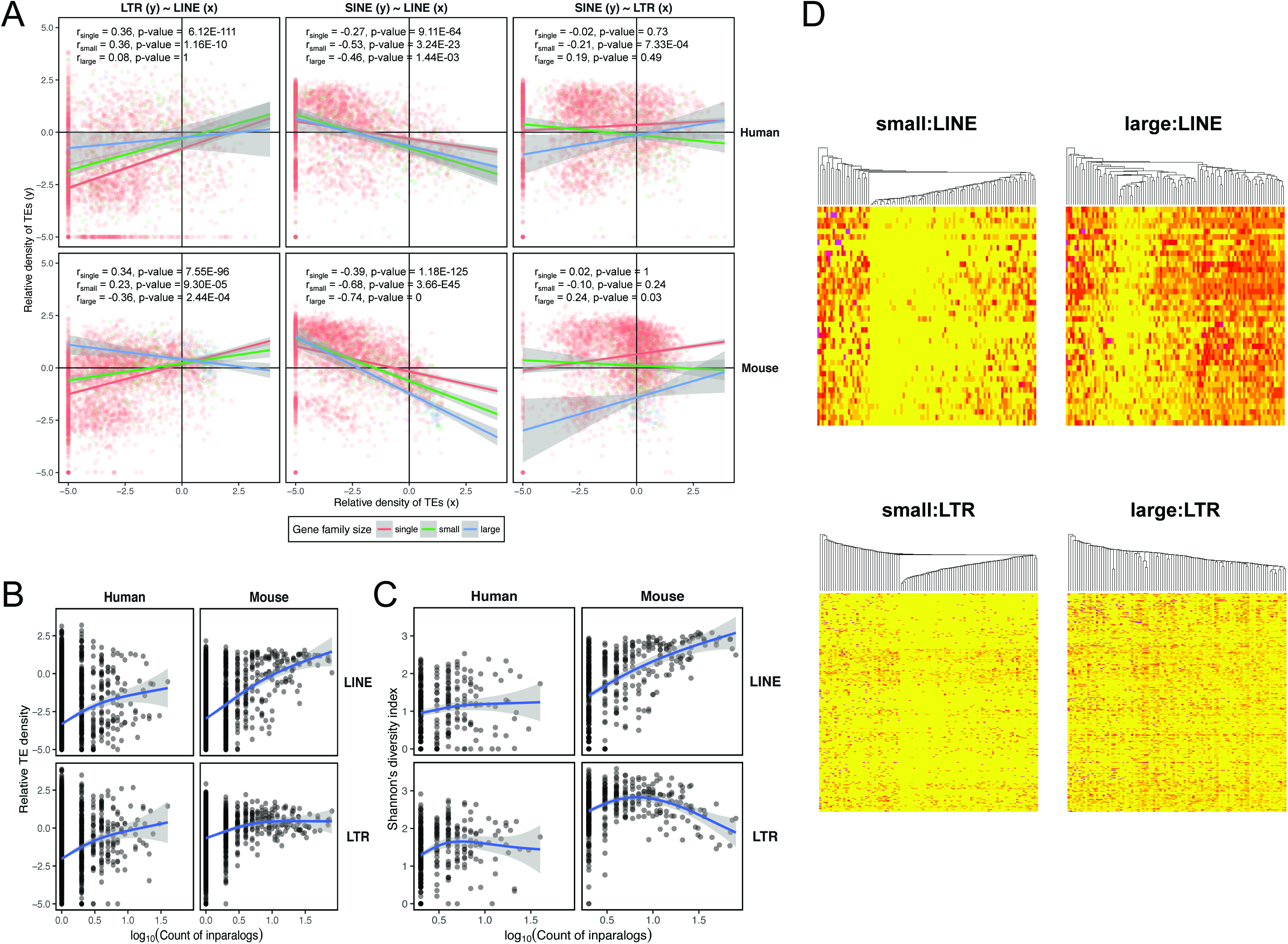
Correlations between gene family size and RT density, diversity and divergence. Correlation of the average lineage-specific RT density between the three RT classes (LINEs, LTRs and SINEs) and the difference in correlation between the three gene family size categories (i.e. single genes, small gene families and large gene families) in the human and mouse genome (A). Correlation was calculated using Spearman’s correlation coefficient. The trend lines and their interactions are based on analysis of covariance (ANCOVA). B) shows the relationship to the gene family size of the lineage-specific RT density, averaged for each gene family. C) shows the relationship between RT subfamily diversity (C) and gene family size. Gene family size is defined as the common logarithm of the count of inparalogs within a gene family and generalized additive models were used to describe the relationship in B and C. D) shows the distribution of RT abundances for the LINE and LTR subfamilies in the mouse genome among the gene families of small and large size. The individual RT subfamilies are represented by rows and gene families by columns in the heatmaps. Yellow corresponds to abundances of zero, and the progressively more intense red colors represent the presence of more elements of particular RT subfamilies. We randomly chose one hundred gene families to visualize RT abundances in both the LINE and LTR panels. The gene families (columns) were hierarchically clustered based on their pattern of abundance of individual RT subfamilies for LINEs and LTRs. RT subfamilies (rows) are ordered according to the average divergence from consensus for a given RT subfamily from the youngest (top) to the oldest (bottom).

**Table 1.** Counts of gene families in the inparalog and outparalog datasets, where the gene family size of one corresponds to the number of families containing only a single gene.

| GF Size | Inparalogs |  | Outparalogs |  |
| --- | --- | --- | --- | --- |
|  | Human | Mouse | Human | Mouse |
| 1 | 3578 | 3541 | 3578 | 3541 |
| 2 | 218 | 225 | 1218 | 1234 |
| 3 | 37 | 46 | 498 | 484 |
| 4 | 41 | 35 | 241 | 243 |
| 5 | 17 | 22 | 141 | 145 |
| 6 | 14 | 13 | 92 | 98 |
| 7 | 11 | 10 | 61 | 60 |
| 8 | 6 | 7 | 40 | 39 |
| 9 | 5 | 11 | 32 | 33 |
| 10 | 0 | 6 | 20 | 15 |
| >10 | 20 | 66 | 124 | 128 |

**Table 2.**
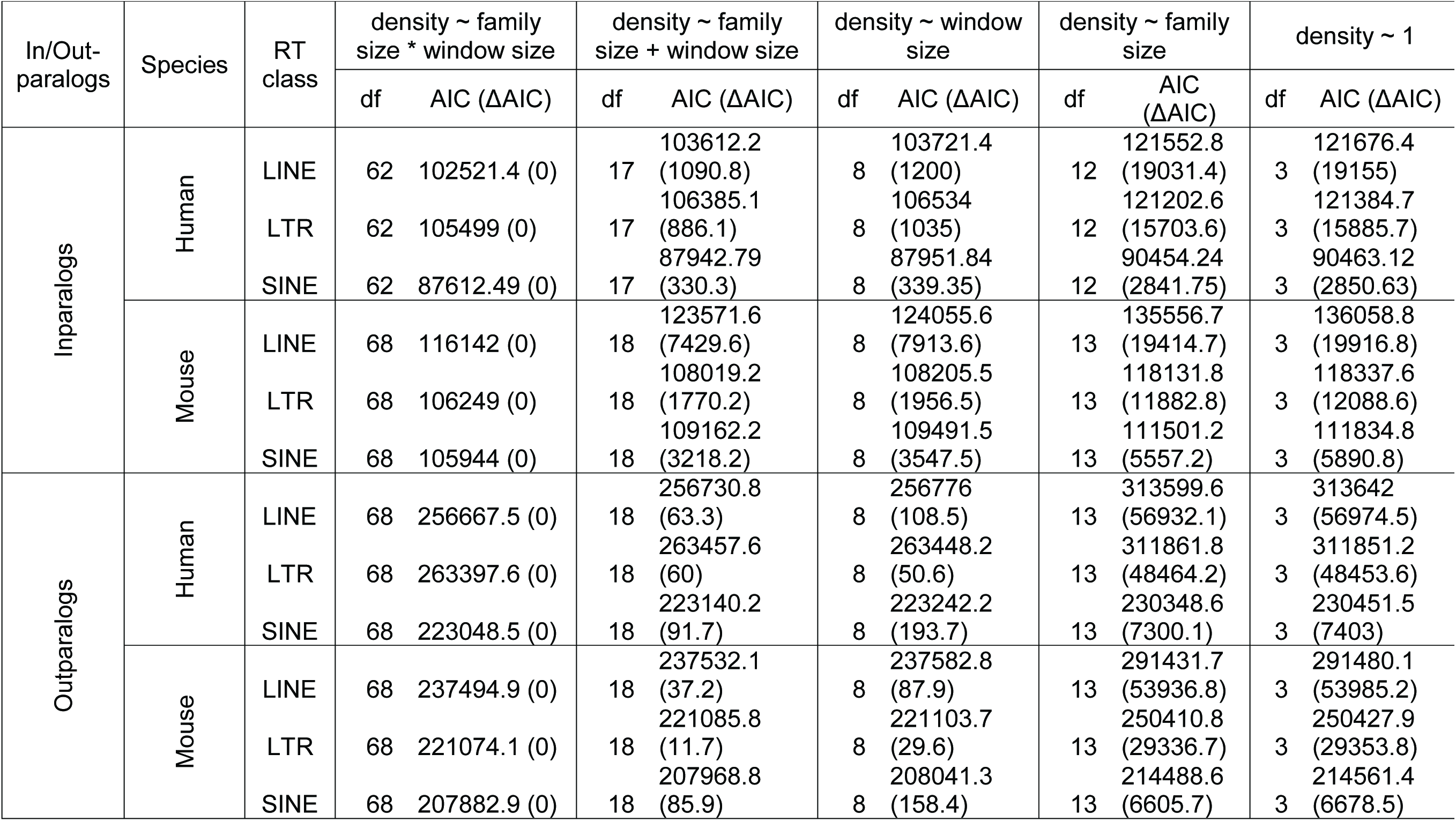
A comparison of models explaining the relationship between gene family size, size of neighborhood (window size) and RT densities, including degrees of freedom (df) and Akaike’s Information Criteria (AIC) values.

In the case of inparalogs, representing lineage-specific gene family expansions, we found a striking increase in the densities of LINEs and LTRs and a decrease in the densities of SINEs, respectively, with increasing gene family size (**Fig. 1A**). This was most pronounced in small windows (10 and 50 Kb), suggesting that RT content is altered in close proximity to genes, whereas, in larger windows (1 and 5 Mb), the RT densities are more similar to the genome-wide average with minimal differences between single genes and large gene families. The LINE and LTR densities for single genes were substantially lower than the genome-wide average, in contrast to dramatically higher RT densities in small gene families (2-5 genes). This suggests that even one or a few lineage-specific duplication events producing a few inparalogs from a single gene have an important effect, especially in the mouse genome.

Despite the fact that the full model fitted the outparalog dataset best (**Fig. 1A**), the patterns of RT densities changed little with respect to gene family size (from one to many genes). There was essentially no difference between single genes and genes of small gene families and only a very slight increase for larger gene families, mostly in the same direction as the densities for inparalogs. The effect of window size for LINEs and LTRs is considerable for gene families composed of outparalogs. The areas close to genes (10 and 50 Kb) have much lower densities of LINEs and LTRs than larger areas (1 and 5 Mb), suggesting that some functional constraints might be occurring. This suggests that the effect of gene family size is important mainly in lineage-specific expansions.

RT density is a function of element abundance and element length and thus it is important to examine the contributions of the two variables separately. The RT abundance is the natural logarithm of RT counts in a given window size and we scaled the counts so that they were comparable between windows of different sizes. We found that the full model, including gene family size and window size, as well as their interaction, was highly significant both for inparalog and outparalog datasets (**Supplemental Table 3**). As in the case of RT densities, the outparalog datasets exhibited much lower log-likelihood ratios than the inparalog datasets. Patterns of change in RT abundance with increasing gene family size closely followed RT density for both inparalogs and outparalogs (**Supplemental Fig. 3**), with low abundances in close proximity (10 and 50 Kb) of single genes and outparalogs and a considerable increase for inparalogs (i.e. lineage-specific expansions). Abundances in larger windows (1 and 5 Mb) around genes were comparable to the genome-wide average for all genes.

**Figure 3.**
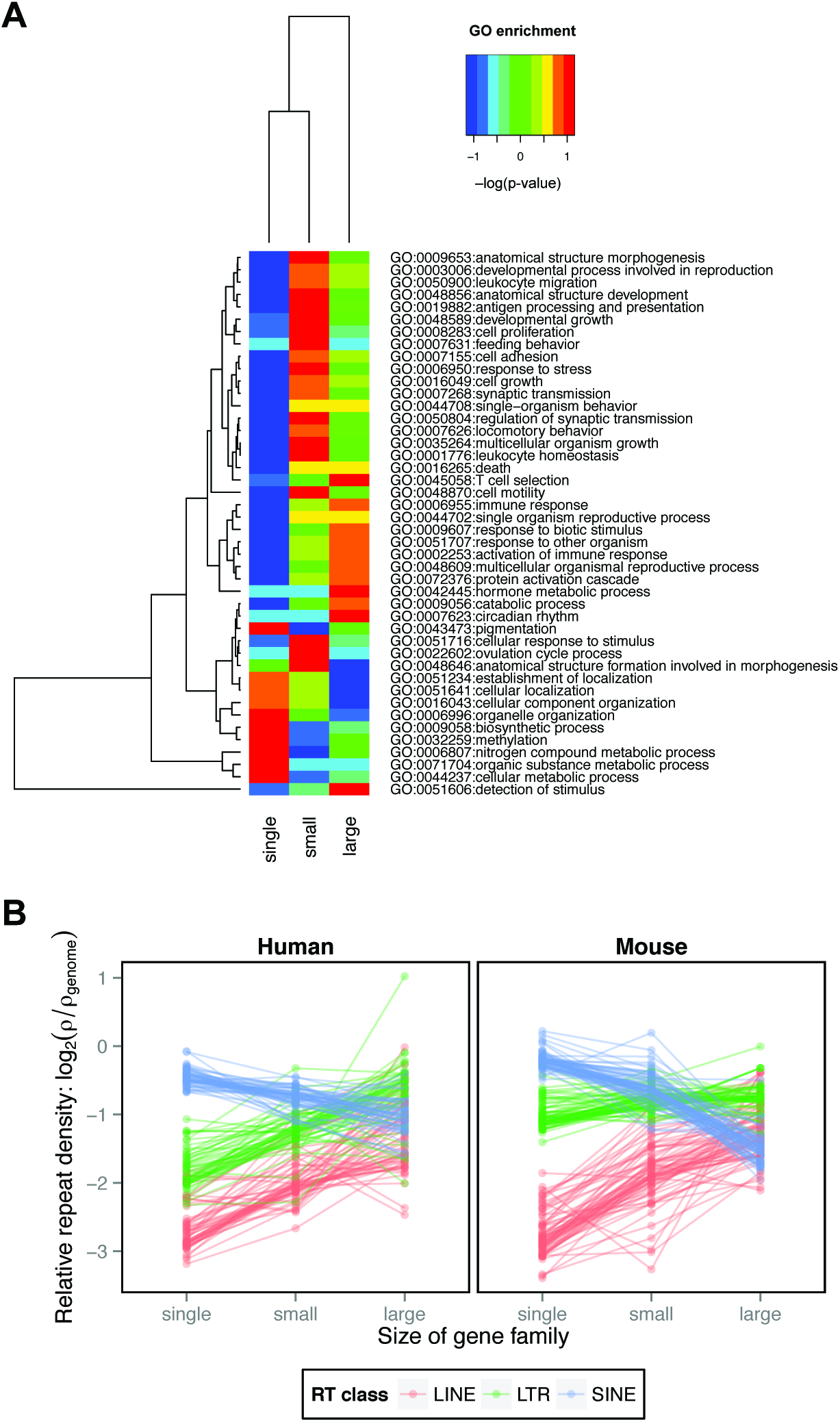
Gene Ontology (GO) data. (A) Examples of functional distinction of gene families of different size (single genes, small gene families and large gene families) based on GO biological processes term enrichment. The warmer the colors the higher the enrichment, while the colder the colors the lower the enrichment. The GO biological processes were clustered based on prevalence between the three gene family size categories. (B) The effect of gene family size (defined as the count of inparalogs) on the average relative density of lineage-specific RTs of the three classes (LINEs, LTRs and SINEs) within the same GO term (all three GO domains included). The average RT densities within the same GO term were connected by lines between the three gene family size categories (i.e. single genes, small gene families and large gene families).

In general, the distribution of average RT length was found to be quite noisy (**Supplemental Fig. 4**). Nonetheless there are clear trends in the distributions of LINEs and LTRs in the mouse genome as both of those RT classes increased in average element size for larger gene families. The pattern was generally highly significant for all inparalog datasets, while the significance for outparalog datasets was weak or absent (**Supplemental Table 4**). This suggests that RT abundance is more important than RT length in driving RT density increase relative to gene family size. However, the average RT element length also contributes to the RT density increase, at least for LINEs and LTRs in the mouse genome.

**Figure 4.**
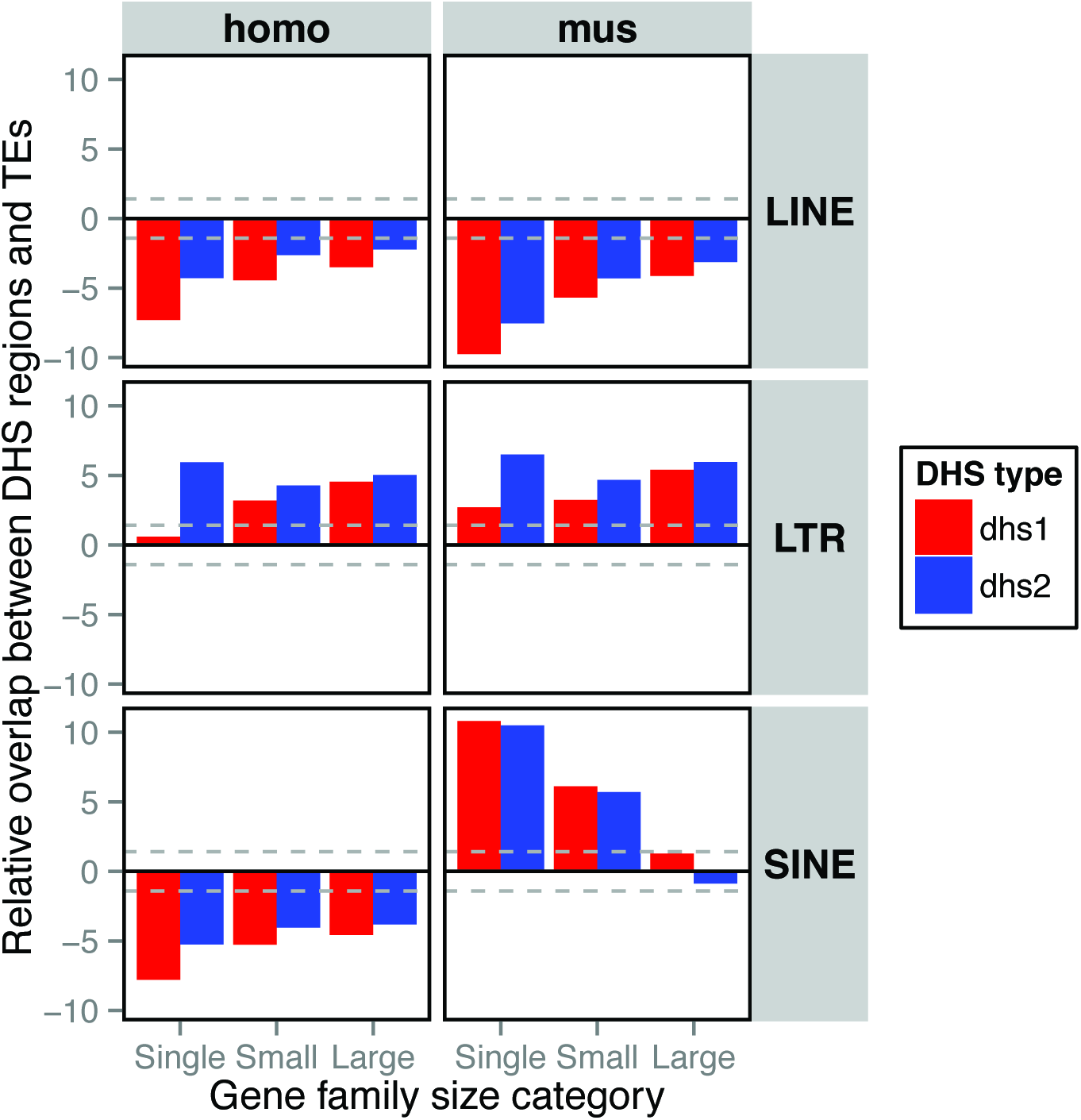
Analysis of the overlap between DNase I Hypersensitive Sites (DHS) and RTs of the three classes (LINEs, LTRs and SINEs) among the three gene family size categories (i.e. single genes, small gene families and large gene families). The overlap between DHS regions and RTs was tested for two sets of DHS regions: the full dataset (DHS1; red) and the dataset of tissue/cell type-specific DHS regions (DHS2; blue). The tissue/cell type-specific DHS regions are those present in ten or fewer tissues/cell types. The positive values indicate that the overlap is higher than the random expectation and the negative values indicate the opposite. The dashed grey lines depict 95% confidence intervals.

We compared the RT content surrounding a representative gene family, *ApoI*, in human and mouse (**Fig. 1B-C** and two other examples are shown in **Supplemental Fig. 5). Figure 1C** shows a phylogeny containing *ApoI* human and mouse genes in different clades with the pig *ApoI* genes in several clades, including one that is a putative outgroup to the human and mouse genes. The *ApoI* regions in the human and mouse genomes are enriched both in LINEs and LTRs but not SINEs. This observation and that of separate clades for the two taxa are consistent with *ApoI* genes being inparalogs in the human and mouse genomes, and thus the gene expansions being lineage specific in the human and mouse genomes.

**Figure 5.**
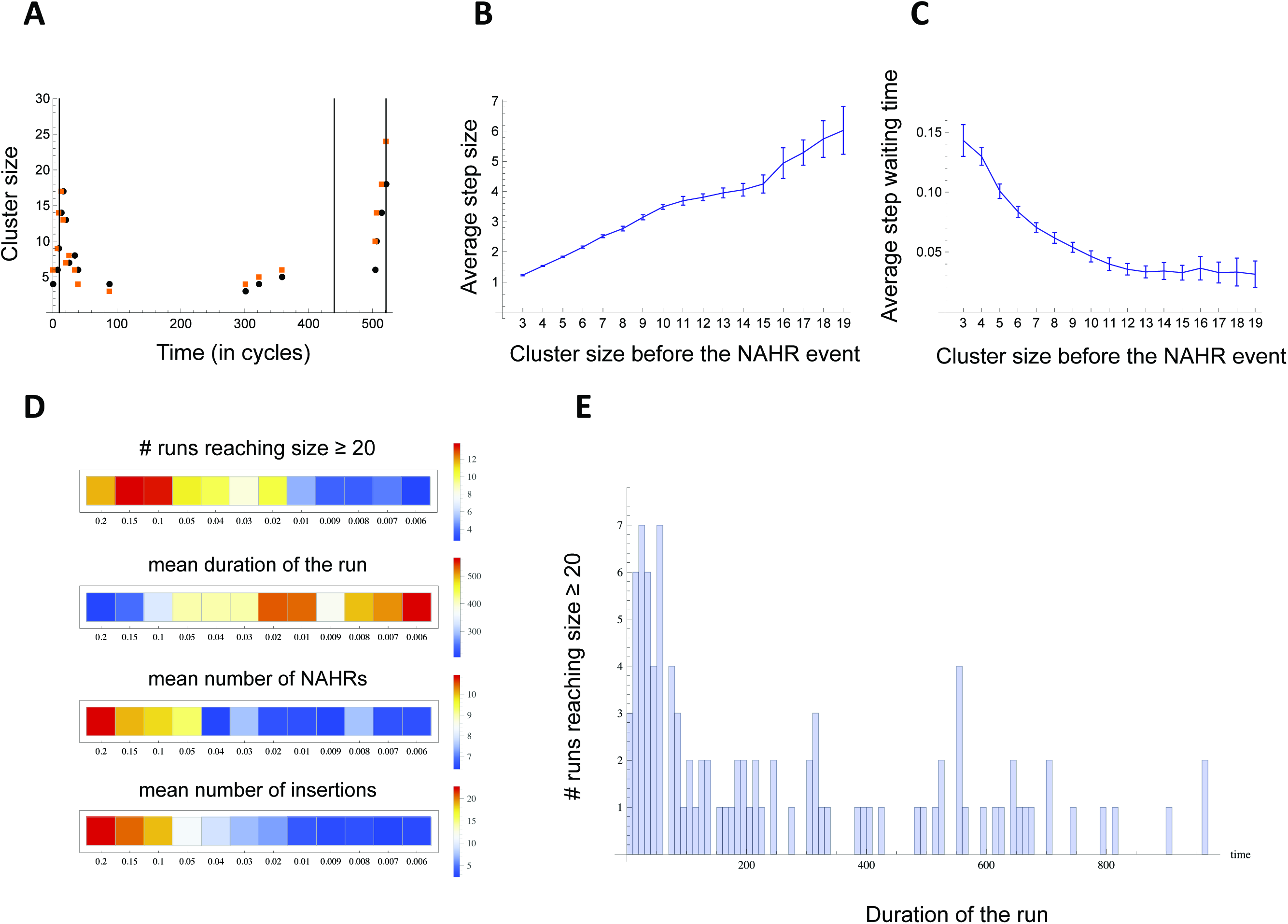
Computer simulation of gene family expansion. (A) An example of a single simulation run that was terminated upon the cluster reaching size = 20. Parameter values were: *ι* = 0.14, μ = 0.4 and *u* = 0.02. Black and orange dots represent the cluster sizes immediately before and after NAHR, respectively. Timings of the fresh LINE insertions are shown by the vertical lines. (B) The average size of gain or loss (i.e. step) in the gene cluster as a result of NAHR for each cluster size category (i.e. immediately before the NAHR occurred). The initial cluster sizes are shown on the X-axis and 95% confidence intervals for the means are indicated by the vertical error bars. (C) The average run duration for each cluster size category. The absolute values of run duration depend on the parameter values chosen, while here we are interested in relative comparison of durations among the cluster size categories. To remove the effects of the parameters, for each combination of the parameter values, the run durations were normalized so that they sum to 1. The data in B and C were pooled over 48,000 runs, with μ varying from 0.1 to 0.4 and *u* varying from 0.006 to 0.2. (D) The effect of the rate of insertion of fresh LINEs, *u*, on (i) the number of “successful” runs that reached size >= 20, shown as the proportion of the 1000 runs, (ii) the average duration of the successful runs, (iii) the mean number of NAHR in the successful runs and (iv) the number of fresh LINE insertions (only the successful runs were considered). (E) An example of the distribution of the times of the last NAHR event among the successful runs. Parameter values were the same as in A. Note the outlier runs where the last NAHR occurred much later then average.

### Correlation between RT densities of the three RT classes and gene family size

We explored the relationship between densities of elements of the three RT classes and asked whether these differ between the three gene family size categories (i.e. single genes, small gene families and large gene families). Employing analysis of covariance (ANCOVA) we identified significant interactions between correlations of RT classes and densities and gene family size (**Table 3**, **Fig. 2A**). Individual correlations were also confirmed by Spearman’s correlation coefficient (Bonferroni correction used: n=3). The SINEs and LINEs exhibited negative correlation, which strengthened as the gene family size grew. Such a negative relationship corresponds to the overall difference in their dependence on GC content for their occurrence (Lander et al. 2001; Waterston et al. 2002). Interestingly, the LTR elements exhibited differences in co-occurrence with the two other RT classes between gene families of the three size categories. Generally, there was a positive relationship between LINE and LTR densities for single genes and small gene families. However, for large gene families this relationship became negative in the mouse and non-significant in the human. More interestingly, the relationship between SINE and LTR densities complemented this finding. There is no correlation between SINEs and LTRs for single genes, a negative correlation for small gene families and a positive correlation for large gene families in both genomes.

**Table 3.** The relationship between densities of the three RT classes with respect to gene family size, where the interaction between the RT density correlation and gene family size was tested using analysis of covariance (ANCOVA).

| Species | Comparison | AIC full | AIC w/o interactions | F | df1,df2 | p-value | slope(single) | slope(small) | slope(large) |
| --- | --- | --- | --- | --- | --- | --- | --- | --- | --- |
| Human | LTR ~ LINE | 16455.58 | 16456.42 | 2.4184 | 2,3884 | 8.92E-02 | 0.37477 | 0.30288 | 0.10097 |
|  | SINE ~ LINE | 14080.67 | 14092.1 | 7.7155 | 2,3884 | 4.53E-04 | -0.16529 | -0.32005 | -0.26106 |
|  | SINE ~ LTR | 14285.28 | 14294.09 | 6.405 | 2,3884 | 1.67E-03 | 0.05405 | -0.10354 | 0.18448 |
| Mouse | LTR ~ LINE | 13552.93 | 13575.18 | 13.148 | 2,3924 | 2.02E-06 | 0.28702 | 0.16341 | -0.13826 |
|  | SINE ~ LINE | 12884.33 | 12908.9 | 14.316 | 2,3924 | 6.38E-07 | -0.24173 | -0.40902 | -0.54104 |
|  | SINE ~ LTR | 13300.52 | 13305.88 | 4.682 | 2,3924 | 9.31E-03 | 0.15512 | -0.05274 | 0.31568 |

### The effect of increasing gene family size on RT density and diversity for large gene families

Given the interaction between LINE and LTR content and gene family size, we also explored how this relationship changes for very large gene families and how it is reflected in the prevalence of individual RT subfamilies. The RT subfamily diversity reflects patterns of RT accumulation. Assuming that only one or a few RT subfamilies of each class are active at any time (Batzer and Deininger 2002), then high RT subfamily diversity reflects high continuous RT accumulation throughout the evolution of a gene family. Comparison of density and diversity for individual RT classes relative to gene family size should reveal patterns of RT deposition over the evolutionary period. We analyzed the relationship between the average RT densities, the overall diversities of RT subfamilies, and the sizes of gene families undergoing lineage-specific expansion (**Fig. 2B, C**). Generalized additive models were used and smoothed relationships were compared to the null model to obtain significances (**Supplemental Table 5**). Our analysis provides contrasting results between the two genomes. In the mouse genome, LINE density and diversity exhibit a steep and continuous increase from single genes to large gene families (density: F_2.37,3926.63_ = 209.96, p-value < 2.2E-16; diversity: F_1.89,429.11_ = 84.483, p-value < 2.2E-16). In the human genome, by contrast, the increase can only be seen for small gene families because the average LINE density and diversity does not change for large human gene families. The significance of the effect of human gene family size in explaining RT diversity is rather weak (density: F_2.01,3886.99_ = 61.587, p-value < 2.2E-16; diversity: F_1.63,309.37_ = 4.4754, p-value = 1.802E-02).

The effect of gene family size is strongly correlated with change in density and diversity of LTRs in both genomes (density_human_: F_1.96,3887.04_ = 51.882, p-value < 2.2E-16; diversity_human_: F_2.53,342.47_ = 9.7455, p-value 1.46E-05; density_mouse_: F_2.53,3926.65_ = 49.004, p-value < 2.2E-16; diversity_mouse_: F_2.83,435.17_ = 19.47, p-value = 2.31E-11). The pattern for LTRs contrasts between RT density and diversity. The LTR density increases for small gene families and stays steady for large gene families in both genomes (**Fig. 2B**). The LTR diversity increases for small gene families, however for large gene families it tends to decrease, especially in the mouse genome (**Fig. 2C**). The contrast in LINE and LTR diversity can be explained by a decrease in deposition of new LTRs and the passive duplication of existing LTRs along with duplicated genes.

### The age of RT subfamilies and their distribution among gene families

We compared distributions of RT abundances for individual LINE and LTR subfamilies for the lineage-specific expansions of gene families of small and large sizes. The LINE and LTR subfamilies were ordered according to the average divergence from consensus from the youngest to the oldest mouse RT subfamilies. The individual gene families were hierarchically clustered according to the abundance pattern among individual RT subfamilies (**Fig. 2D**; the full data for both species are in **Supplemental Fig. 6**). It is clear that more LINE and LTR subfamilies are represented in the regions surrounding larger gene families, however there is no apparent age preference in the RT subfamilies surrounding any size gene family because all ages of subfamilies are found in the gene family regions. Thus there appears to be essentially no distinction for gene family size based on the presence of elements of specific RT subfamilies.

The detailed patterns are also interesting because they differ somewhat between LINEs and LTRs. For small and large gene families, one can find gene families lacking essentially any LINEs, gene families over-populated by elements of essentially all LINE subfamilies and those which represent a transition between these two extremes. On the contrary, LTRs populate moderately almost all gene families without distinction to gene family size. As for the variability in abundance of RT subfamilies within individual gene families, there is little variability for LINE subfamilies within any gene size category. Individual LTR subfamilies vary greatly in their abundance, especially for large gene families with some subfamilies contributing many elements while others contribute none (**Fig. 2D**). Specific differences in the activity of RT subfamilies between the two RT classes may cause such a distinction, however it is unlikely to explain differences in variability between individual gene families.

### Correlation of RT content in homologous gene families between the human and mouse genomes

Co-occurrence of elements of different RT subfamilies associated with the same gene family led us to explore interspecies correlation of RT densities for the homologous gene families. We found a highly significant correlation between the average RT densities for individual homologous gene families between the human and mouse genomes (**Supplemental Table 6**). The RT densities represent lineage-specific elements, but nonetheless their occurrence is correlated between the two genomes. This suggests that there was a common history in the environments of the homologous genes which allowed the accumulation of RT elements in the human and mouse lineages. These correlations are stronger for LINEs and SINEs (r^2^_human LINE single_= 0.23, r^2^_human LINE small_ = 0.31, r^2^_human LINE large_ = 0.28, r^2^_mouse LINE single_ = 0.25, r^2^_mouse LINE small_ = 0.27, r^2^_mouse LINE large_ = 0.14, r^2^_human SINE single_ = 0.41, r^2^_human SINE small_ = 0.32, r^2^_human SINE large_ = 0.27, r^2^_mouse SINE single_ = 0.41, r^2^_mouse SINE small_ = 0.31, r^2^_mouse SINE large_ = 0.16) than for LTRs (r^2^_human LTR single_ = 0.10, r^2^_human LTR small_ = 0.07, r^2^_human LTR large_ = 0.17, r^2^_mouse LTR single_ = 0.11, r^2^_mouse LTR small_ = 0.03, r^2^_mouse LTR large_ = 0.07). This suggests that the common/shared history of homologous regions between human and mouse is more important for LINEs and SINEs than for LTRs, which tend to vary more in density between homologous gene families of the two genomes. LTRs thus seem to be less dependent on genomic region history than the other two RT classes.

### RT densities within the same functional class

Large gene families were previously shown to be enriched for specific functions contrasting with the functional composition of single genes (Emes et al. 2003). We confirmed this for our dataset by conducting GO term enrichment analysis among gene families of the three size categories (**Fig. 3**, **Supplemental Fig. 7**). Small gene families are more highly associated than average (red) with about half the GO term functions, where singles genes have a lower-than-average association (blue) and large gene families have for the most part an average association (green). Single genes are associated more than average with a smaller percentage (~20-25%) of GO term functions and large gene families are associated also with about ~20-25% of functions, but these are different ones than the single genes (blue) and the small gene families (green). There are categories of GO terms that are enriched in large gene families (red) and are less often associated with single genes (blue), and vice versa. When this is the case, the small gene families are intermediate in enrichment (green). There are also many GO categories enriched in small gene families (red), neutral in large families (green), and less commonly assigned to single genes (blue). Single gene families show associations with GO terms that are essentially opposite those of small and large gene families. Also, there is some functional distinction between small and large gene families.

Had the RT accumulation been associated with a specific function, such a functional distinction between gene families of the three sizes might have been responsible for differences in the RT prevalence we observe for gene families of different size. To test whether the actual size of the gene family rather than gene function reflects the altered RT content, we studied average repeat densities as a function of gene family size associated with the same GO term (**Fig. 3B**). We found increases in LINE and LTR densities for all of the GO terms between single genes, small and large gene families. By contrast, the SINE densities exhibited the opposite pattern. All comparisons were highly significant (**Table 4**). Thus, the altered RT content appears to be characteristic for the multi-copy nature of gene families without respect to the functional categories into which the genes fall. However, the effect of function cannot be discounted completely.

**Table 4.** Analysis of the effect of gene family size on explaining RT densities when included only gene families associated with a GO term present in all three gene family size categories, using the generalized Least Squares (GLS) model to capture correlation structure between individual GO terms.

| Species | RT class | AIC (GF size) | AIC (null) | LogLikelihood Ratio | P-value | RT density estimate |  |  |
| --- | --- | --- | --- | --- | --- | --- | --- | --- |
|  |  |  |  |  |  | single | small | large |
| Human | LINE | -902243.2 | -901374.5 | 872.7 | < 0.0001 | -2.78 | -1.87 | -1.39 |
|  | LTR | -898863.5 | -898250.4 | 617.2 | < 0.0001 | -1.95 | -1.17 | -0.64 |
|  | SINE | -919759 | -919634.5 | 128.5 | < 0.0001 | -0.53 | -0.76 | -1.03 |
| Mouse | LINE | -839461 | -838015.9 | 1449.1 | < 0.0001 | -2.82 | -1.89 | -1.05 |
|  | LTR | -847997.3 | -847833.6 | 167.8 | < 0.0001 | -1.12 | -0.77 | -0.89 |
|  | SINE | -863464.5 | -861751.5 | 1717 | < 0.0001 | -0.24 | -0.60 | -1.61 |

### Association of RTs and open chromatin regions

The Encyclopedia of DNA Elements (ENCODE; (Consortium 2012; Stamatoyannopoulos et al. 2012)) catalogs DNase I Hypersensitive Sites (DHS) data representing regions of open chromatin and thus regulatory activity in genomes. We used DHS data from the human and mouse genomes to test the hypothesis that a higher abundance of RTs is associated with regulatory activity of genes in recently expanded gene families. We assessed potential overlap between the three classes of RTs and the DHS around genes of the three gene family size categories and found occurrences that were significantly higher or lower than the null expectation of randomly distributed DHS (**Fig. 4**).

DHS significantly overlap LTRs around single genes and both small and large recently-expanded gene families, supporting the regulatory hypothesis. There were, however, conflicting results between the two sets of DHS regions, DHS1 and DHS2. While we found an increase in the association of LTRs and DHS with increased gene family size for the whole DHS dataset (DHS1), the cell type/tissue-specific DHS2 exhibited the highest association for single genes with a drop in small gene families and only a slight increase for large families. In contrast to LTRs, LINEs were consistently underrepresented in the DHS regions in both genomes. SINEs, on the other hand, exhibited conflicting results, being systematically underrepresented in the human genome and significantly enriched around single genes and small gene families in the mouse genome. The trend in the mouse genome was a decrease in the association of RTs and DHS regions with increasing size of gene families. This strong contrast for the SINE pattern between the human and mouse lineages is similar to enrichment of regulatory factor CTCF-binding sites in lineage-specific SINEs found in rodents but not in primates and humans (Schmidt et al. 2012).

### Simulation of a second phase of rapid gene family expansion

Our observation that the size of a gene family is an important predictor of the RT distribution in close proximity to the family members suggests the direct involvement of the RTs in the expansion process. This could have been achieved through LINEs serving as the homologous sequences for NAHR in what we suggest is a second phase of gene family expansion characterized by rapid gene duplication due to continuous accumulation of LINEs (see Discussion). Intuitively, the gene clusters with duplications resulting from NAHR could be prone to further NAHR events with progressively more opportunities of forming additional homologous sequences. We feel that this process could even take on runaway proportions. However, decay in homology between LINEs may decrease the probability of NAHR and slow down the whole process of expansion. Using a simulation, we qualitatively assessed the conditions potentially leading to a runaway gene expansion process as defined below.

We developed a model to test our hypothesis that, in a second phase of gene family expansion, continuous accumulation of RTs causes rapid gene duplication (the simulation algorithm is illustrated in **Supplemental Fig. 8**). The population, initially fixed for a particular small gene family genotype, is sampled at discrete time steps. The time between steps is assumed to be sufficiently long for the mutant type to reach fixation (i.e. ~4Ne generations in the neutral case). At each step, the resident genotype either stays or, with the fixation probability *ι* (iota), is replaced by a new genotype and subjected to a new insertion of LINEs (with probability *u*) and to NAHR (with the probability *μ*). Due to the theoretical complications of calculating the exact fixation probabilities in large populations (Weissman and Barton 2012), the parameter *ι* is set arbitrarily. However, it is assumed to be dependent on the rate of recombination, the effective population size and/or any form of selection altering local fixation probabilities.

The simulation was run for a range of values of *ι*, *u* and μ, for either 1,000 or 50,000 repetitions for each combination of the three parameter values. Each run was stopped in one of two cases: (i) the number of genes in the cluster reached 20 or (ii) the number of simulation cycles reached a threshold of 1,000/50,000. In a typical run (**Fig. 5A**) most of the gene family size change occurred during relatively short time periods at the very beginning (not quite reaching the size threshold of 20) and near the end when the gene family size threshold was reached and the run stopped. **Figures 5B** and **C** illustrate the two key properties of the cluster size dynamics for the combined results of 48,000 runs over a wide range of parameter values. First, the larger clusters tend to undergo larger duplications or deletions during NAHR, and this effect is independent of the simulation parameters even though the variability in the size of duplication/deletion increases as the size of the cluster increases (**Fig. 5B**). Second, the average duration of runs is shorter in the large clusters relative to the small clusters (**Fig. 5C**). In short, the cluster size dynamics speed up (in either direction) as the gene cluster grows in size. Without an upper boundary this process has the potential to quickly create a very large number of genes, hence our choice of the term “runaway expansion”. This led us to seek an explanation for the apparent shift in the behavior of gene expansions that appears in **Fig. 5 B** and **C** to occur at a cluster size of ~10. We provisionally define those gene expansion events in which the average step size continues to rise and average run duration time continues to diminish (**Fig. 5 B** and **C**) as having taken on the characteristics of a runaway expansion. By contrast, the other events that show the average step size leveling off (or dropping) and average run duration leveling off (and sometimes even rising) have escaped the runaway expansion fate. As one would predict, these different tendencies create a noticeable increase in variance for both curves **Fig. 5 B** and **C** show.

Increasing parameter values for *ι*, *u* and μ increased the proportion of runs that reached the size threshold criteria of 20 genes (endpoint, **Supplemental Figs. 9A-11A**). In those runs that reached the endpoint size criteria of 20 genes, increasing all three parameters decreased the time to reach the threshold (i.e. the duration of the run, **Supplemental Figs. 9B-11B**), and correspondingly, increased the mean number of NAHR events during the run (**Supplemental Figs. 9C-11C**). At the same time, decreasing the rate of NAHR (μ) allowed for the accumulation of more fresh LINEs, for a constant rate of insertion, *u* (**Supplemental Figs. 10D, 11D**), whereas increasing *u* had an expectedly positive effect on the frequency of insertions (**Fig. 5D, Supplemental Figs. 9D, 10D**). A certain degree of irregularity of the average response to the parameter variation, most notably in the average duration of the run (**Supplemental Figs. 9B-11B**), can be explained by the fact that the distributions of times were often positively skewed, with a considerable proportion of the simulation runs taking much longer to reach the threshold size > 20 for the same combination of parameter values. **Figure 5E** provides a single example of such a distribution with the main mode around 40 and a flat right tail, and with occasional smaller modes around 450 and 900 time cycles (**Fig. 5E**).

Our model did not expressly include selection that would favor an increase in gene copy number, however, keeping all other things constant (rate of recombination, effective population size), *ι* can be interpreted as the fixation probability due to positive selection. **Figure 6** depicts a change in the summary statistics over various values of *ι* showing that positive selection (i.e. higher *ι*) can speed-up the process of gene family expansion by reducing the average duration of a run. More interestingly, increasing the fixation probability due to positive selection may lead to a locally increased density of LINEs even when the rate of a new LINE insertion is constant. Such an outcome can explain the high densities of LINEs we observe in the human and especially in the mouse genome.

**Figure 6.**
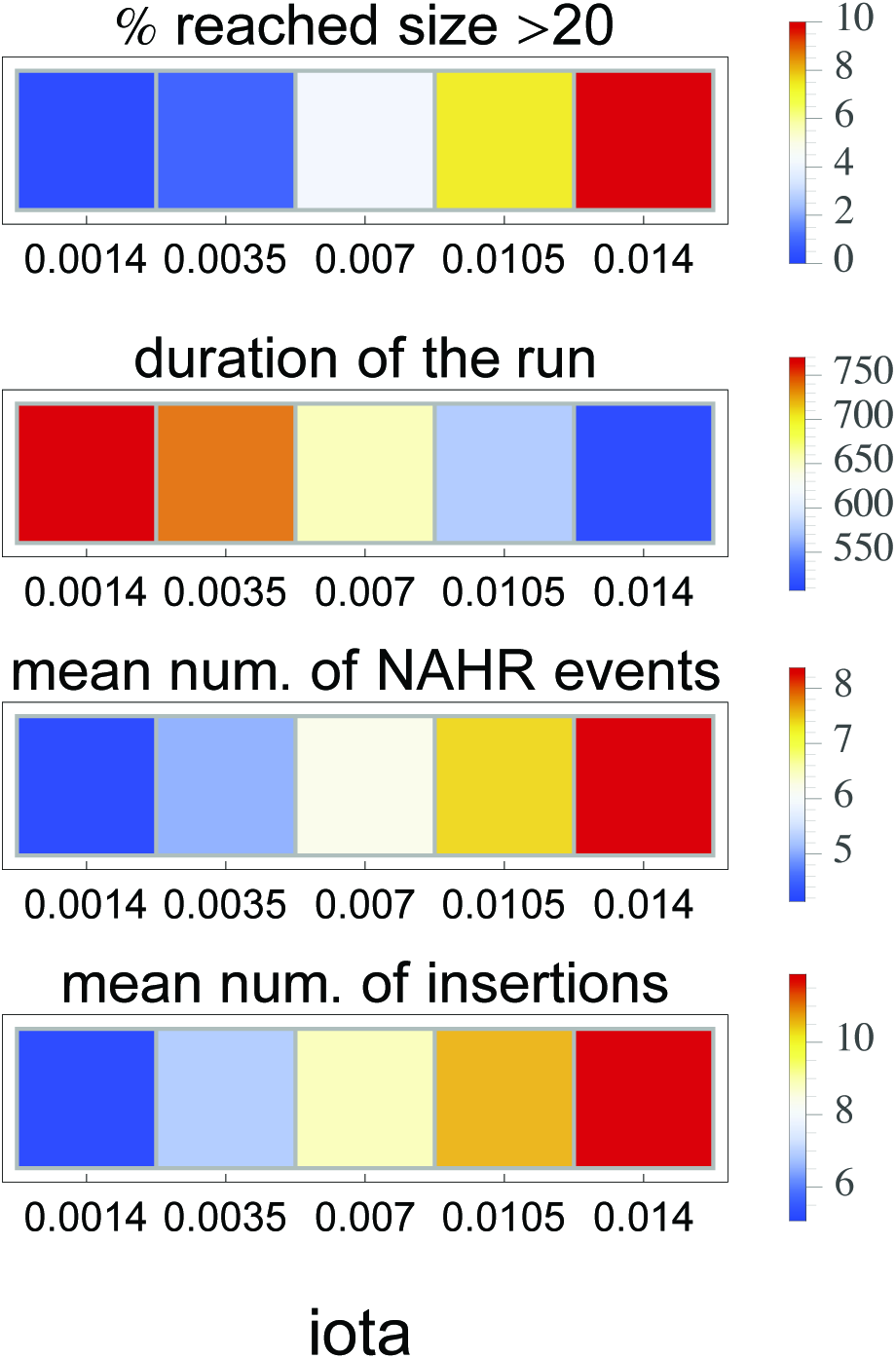
The effect of varying the iota parameter on the progress of a runaway process. In general, the iota parameter reflects processes such as genetic drift, positive selection and/or the rate of recombination.

## DISCUSSION

Retrotransposons have been recognized recently as important forces shaping mammalian genomes and contributing considerably to organismal complexity. In a previous study (Janoušek et al. 2013), we found that LINE and LTR retrotransposons contributed to the evolution of the *Androgen-binding protein* (*Abp*) gene family in the mouse genome. Here we extend that study on a genome-wide scale to the human and the mouse, focusing on the general importance of these two RT classes in gene family expansion. We provide evidence that LINEs and LTRs interact in that process and we suggest that at least some gene family expansions can be divided into two phases. In this hypothesis, the first phase represents sub/neofunctionalization associated with rewiring regulatory networks by LTR elements, while the second phase accelerates gene family expansion by the continuous accumulation of LINEs, and in some cases this process may reach runaway proportions. We support this mechanism with a computer simulation.

### LINEs and LTRs are associated with gene family expansions

We contrasted RT content between lineage-specific and ancestral gene family expansions and found a striking association between RTs and gene family size of lineage-specific expansions in both the human and mouse genomes. This contrasts with only a weak or nonexistent relationship between RT content and ancestral gene families that were fully expanded in the common ancestor of mice and humans, suggesting that the altered RT content is directly related to the expansion process. There are low densities of LINEs and LTRs around single genes with a subsequent increase as the size of gene families expands in both genomes, whereas SINE content differs between the human and mouse genomes. The effect was also strong when RT densities were compared within the same functional category, thereby discounting functional specificity in driving RT accumulation. This emphasizes the role of the duplication/expansion process as such rather than functional differences between single genes and expanded gene families. LINEs and LTRs are enriched at the edges of segmental duplications (a mechanism leading to gene family expansion) in the mouse genome (She et al. 2008) and these elements provide a substrate for NAHR (Campbell et al. 2014; Startek et al. 2015). LINE elements have been found to be a substrate for NAHR producing the most recent duplications in the *Abp* gene family region (Janoušek et al. 2013). Bailey (Bailey et al. 2003) suggested that *Alu*s (SINEs) are the elements enriched at the edges of segmental duplications in the human genome. This contrasts with our data, however, Kim (Kim et al. 2008) proposed that SINEs and LINEs might have been involved in producing segmental duplications at different times in the evolution of primates and that this could have been related to the different activity of these two RT classes. *Alu*s might have played this role during their activity burst ~ 40 MYR ago, whereas LINEs may be a more recent source of structural changes (Kim et al. 2008). This differential timing might have obscured the pattern of SINE densities around expanded gene families in the human lineage. The fact that we see no strong relationship between SINEs and gene family size in the mouse lineage might be related to specific differences of SINE elements between the human and mouse lineages (*Alu*s are ~300 bp in the human lineage vs. ~150 bp SINEs in the mouse lineage). Nevertheless, the signal provided by LINEs and LTRs is strong in both lineages. The question remains, however, whether their structural role is adequate to explain the higher densities and counts associated with lineage-specific gene family expansions.

### Distinct roles of LINEs and LTRs in gene family expansions

Another of our important findings was an interaction between LTRs and the other two RT classes with respect to gene family size, an observation we have not found in earlier reports. LTRs seem to co-occur with LINEs around gene families of small size, however as the gene family size grows, the density of these two RT classes begins to correlate negatively. This is supported by a relationship between LTRs and SINEs. In the mouse genome, the continuous increase in LINE density and diversity in large gene families contrasts strongly with the unchanging density and decreasing diversity of LTRs. Such a pattern can be explained by a decrease in deposition of new LTRs and the passive duplication of existing LTRs along with LINE-dependent duplication of genes in large mouse gene families. This contrasts with the human genome where the densities and diversities of both RT classes (LINEs and LTRs) stay the same for large gene families.

The role of LTRs in this context is understandable because RTs have been shown to contain binding sites for transcription factors (Jordan et al. 2003; Polavarapu et al. 2008). The LTR class has been identified as the main contributor to open chromatin regions and transcription factor binding sites (Jacques et al. 2013; Sundaram et al. 2014), and elements of the LTR class are recruited as tissue-specific promoters by the *Neuronal apoptosis inhibitory protein* (*NAIP*) gene family (Romanish et al. 2007). We found a highly significant overlap of LTRs with DHSs around genes of gene families, which supports their role in gene regulation and suggests that LTRs might play a role in subfunctionalisation of newly duplicated genes. By contrast, LINEs exhibited less overlap with DHS than expected by chance, ruling out their potential involvement in evolution of regulation during gene family expansions.

### Two phases of gene family expansion: A model for retrotransposon interaction

Karn and Laukaitis (Karn and Laukaitis 2009) proposed that the mouse *Androgen-binding protein* (*Abp*) gene family expanded in two phases. First, single genes duplicated to produce two daughter genes in inverse adjacent order (Katju and Lynch 2003). Later, blocks of genes duplicated by NAHR resulting in new genes in direct adjacent order and accelerating the expansion of the ABP gene family. The presence of ERVII (LTR) and L1 (LINE) repeat families in high densities in the mouse and rat *Abp* gene regions with corresponding depletion of other families suggested a functional role in duplication (Janoušek et al. 2013). While ERVII subfamilies were distributed approximately equally between lineage-specific and lineage-shared subfamilies in both genomes, the majority of LINE1 repeats (>90% in the mouse and >80% in the rat genome) were lineage-specific. Thus, >50% of ERVII repeat content originated from insertions that occurred near the ancestor of the *Abp* gene family, while almost no LINE1s were present in the *Abp* region before its expansion. Finally, (Janoušek et al. 2013) identified the break point in members of a LINE1 (L1Md_T) retrotransposon that caused the last NAHR-mediated duplication of the block of genes in the center of the *Abp* region in the mouse genome.

The gene family expansion study we report here may provide a mechanism for the *Abp* gene expansion model proposed by Karn (Karn et al. 2014). They observed that the *Abp* paralogs expressed in the lacrimal and salivary glands are found in different ancestral *Abp* clades and found instances of extremely low levels of paralog transcription without corresponding protein production in one gland with high expression in the other. They proposed a model in which genes expressed highly in both glands ancestrally were down-regulated subsequent to duplication as the result of subfunctionalisation, and they suggested that the most parsimonious point for this would be when the first <*Abpa-Abpbg*> gene module duplicated to produce a pair of daughter modules in inverse adjacent order (Karn and Laukaitis 2009; Karn et al. 2014). We think that this process involved the accumulation of ERVII retrotransposons (Janoušek et al. 2013), which are LTRs that could have ultimately been responsible for the subfunctionalisation of the daughter <*Abpa-Abpbg*> gene modules by modifying gene regulation.

This is consistent with the ERVIIs accumulating in the mouse and rat gene regions before the LINEs (Janoušek et al. 2013) and our finding in this report that small gene families exhibit an increase in density and diversity of LTR elements in both genomes along with increasing size of lineage-specific gene family expansion. Because LTRs have been suggested to play a role in regulation of gene transcription (Feschotte 2008; Bourque 2009), we propose that their accumulation is important in a first phase of gene expansion characterized by sub-/neo-functionalisation (Lynch and Force 2000). The second phase of gene family expansion is characterized by continuous deposition of LINEs, which may make the whole region more volatile, promote further rapid expansion, and decrease the rate of LTR accumulation. The subsequent lineage-specific accumulation of the LINEs in the *Abp* gene region (Janoušek et al. 2013) is consistent with the notion that they mediated the second phase of *Abp* gene expansion by NAHR.

Finally, we suggest that the mouse *Abp* gene family might represent an example of runaway gene expansion. Karn and Laukaitis (Karn and Laukaitis 2009) argued that the second phase of mouse *Abp* gene expansion occurred in the largest and most volatile *Abp* central clade where blocks of multiple *Abp* gene modules were duplicated by NAHR, pushing the ancestral gene sets apart and leaving the more diverged sequences on the flanks. NAHR accelerated this process dramatically, characteristic of the “snowball effect” of (Kondrashov and Kondrashov 2006), not to be confused with ‘snowballing’ in speciation (Orr 1995). Indeed, 36% of extant *Abp* genes were formed by the last two large block duplications. One consequence of this was volatility evidenced by copy number variation (CNV) in this central clade and extensive duplication of pseudogenes (i.e. quadrupling the number of pseudogenes in the final two duplications) along with potentially expressible genes (Janoušek et al. 2013; Karn et al. 2014).

### A computer simulation supports our second phase of gene family expansion

The snowball effect (Kondrashov and Kondrashov 2006) amounts to a rapid, local increase in gene duplications caused by a high rate of LINE accumulation, subsequent NAHR and, possibly the presence of low copy repeats (LCRs) produced by previous duplications (Karn and Laukaitis 2009). Given the continuous accumulation of LINEs serving along with the LCRs as break points for NAHR, the whole region can expand rapidly. If it becomes extensive enough, this second phase could be described as a runaway process.

To explore this possibility, we conducted a computer simulation that showed that rapid increase in gene copy number is a possible outcome of increased deposition of homologous sequences representing LINEs and that, in some cases, the gene expansion may reach runaway proportions. This was true even though the simulation accounted for the fact that previously incorporated LINEs undergo loss of their homology over time due to mutation. We found that the number of gene copies created in one NAHR event increased with increasing size of the gene cluster and, more importantly, that the time between two consecutive NAHR events is shorter for larger gene clusters. Both of these findings correspond to speeding up gene family expansion and formation of large blocks, i.e. LCRs, spanning several genes. Interestingly, this process is possible even without invoking selection in favor of increased gene copy number; however, this does not preclude the possible role of selection for increased gene dosage. In our simulation, we did not specifically test the direct involvement of selection favoring the increase in gene copy number. Nevertheless, the fixation rate of the new genotype (*ι*) will increase whenever the recombinant genotypes are favored, thus reducing the duration of each run. Besides increasing the number of runaway events in our simulations, such an increase in *ι* also leads to the reduction of the total time necessary for an expansion and to an increase in the average number of repeats that accumulated around the genes.

Because all parameters in our model were set arbitrarily, it is difficult to directly relate the quantitative results of the simulations to real-world population genetic data on the human or the mouse. Direct use of this information, although desirable, is methodologically challenging due to the complication of the population genetics theory in large populations (Weissman and Barton 2012). One quantitative characteristic of our simulation that can be interpreted to a certain degree is the evolutionary time required for the runaway process to occur. The duration of the time unit in our model is assumed to be sufficient for a replacement genotype to reach fixation and this time can be reduced or increased drastically in the case of positive or negative selection, respectively, acting on the gene cluster. Given the rather large estimates for N_e_ in the mouse and human, we conclude that a runaway expansion under negative selection is unlikely. Even under neutral evolutionary conditions, a large part of the parameter range we investigated can take a considerable amount of time. For example, with N_e_ = 50,000 and 2 generations per year in the mouse, 400 cycles translates into 40 MYR, which is still less the ~80 MYR since the mouse-human split (Hallstrom and Janke 2010). Invoking selection for increased copy number might therefore be necessary to explain the gene family expansions in both taxa.

### A runaway process in the human and mouse lineage

One of the criteria for entering the second phase could be functional constraint imposed on genes. For instance, (Korbel et al. 2008) found successfully duplicated genes have been described as being located at the periphery of protein interaction networks. About 10% of genes are found to be highly volatile and subject to frequent duplication, deletion and pseudogene formation (Lander et al. 2001; Waterston et al. 2002; Gibbs et al. 2004). These generally possess functions including chemosensation, reproduction, host defense and immunity, and toxin metabolism. We also found distinct GO term enrichment among gene families of different sizes. Genes in this set that are expanded in one lineage are often expanded in another, suggesting similar functional and/or structural pressures (Ponting and Goodstadt 2009). However, we found contrasting patterns of LINE and LTR accumulation between the human and the mouse.

Analysis of recent segmental duplications between human and mouse genomes highlights interesting differences in the distribution of recent duplications (She et al. 2008). Duplications in the mouse genome occur in discrete clusters of tandem duplications, whereas duplications in the human tend to be scattered across the genome. The scattered nature of expanded gene families in the human genome may decrease the chance for gene families to enter the second phase, where the tandemly-arrayed nature of gene families in the mouse may facilitate the runaway process in some cases. Although the percentage of recent segmental duplications between the two genomes is similar (She et al. 2008), larger families are more frequent in the mouse genome (**Table 1**), suggesting that the mouse lineage is richer for gene family expansion events. In addition to specific environmental challenges this lineage might have faced during its evolution, ~20x higher activity of LINEs in the mouse genome (Goodier et al. 2001; Brouha et al. 2003) could have contributed to the more massive expansions in this lineage. Assuming that this figure corresponds linearly into the 20-fold difference in the rate of insertion, *u*, in our simulation results, it seems to us that the difference in the LINE activity between the human and mouse genomes alone might have profound effects on the frequency of the runaway process in the expansion of gene families. Besides functional constraints, the specific nature of duplications and overall activity of LINEs in a given lineage could be important in facilitating the runaway process.

### Common history and duplicability

RTs of the same class accumulate in homologous genomic regions in the human and mouse lineages (Yang et al. 2004). Here, we report the correlation of lineage-specific RT densities between human-mouse homologous gene families that contain lineage-specific expansions (**Supplemental Table 6**). Furthermore, analyzing patterns of accumulation of elements of individual subfamilies revealed that gene families of high RT density exhibit high prevalence of all ages of elements. This was most notable for the LINE class, with strong correlation in the human-mouse comparison as opposed to LTRs. Weak or no correlation for LTRs may correspond to their lineage-specific accumulation depending more on the adaptive needs of a given lineage than on a region-specific feature shared between lineages.

Duplicability is a measure of the likelihood of gene duplication during evolution, which is the product of the rate of mutation producing duplicate genes and the probability that the duplicates are fixed and retained in the genome (He and Zhang 2005; Qian and Zhang 2008). Given their putative involvement in structural changes, LINEs may be associated with duplicability of genomic regions. Such features may be shared between related lineages and cause some gene families to undergo independent expansions in different species.

## METHODS

### Gene family data

We used Ensembl’s Core and Compara databases (Version 75; (Vilella et al. 2009) to obtain human (*Homo sapiens*), mouse (*Mus musculus*) and pig (*Sus scrofa*) genes from their genomes and to determine their evolutionary relationships. The protein coding status of human and mouse genes was confirmed using HUGO/MGI databases (Povey et al. 2001; Blake et al. 2014) and all pseudogenes were discarded from our dataset. The Ensembl Perl API interface was used to obtain the data. We created a pipeline using in-house perl/shell scripts; see details in **Supplemental Fig. 1**. First, we defined clusters of homologous genes of the three species and aligned them using eight alignment tools and obtained consensus protein alignment using M-Coffee (Wallace et al. 2006; Moretti et al. 2007). The low scoring parts of alignments were discarded. We used the TREEBEST program to construct gene family trees for the three species based on aligned clusters of homologous genes and extracted human and mouse lineage-specific duplication events. The pig genes served as an outgroup. The approach is described in detail in (Vilella et al. 2009). Based on this procedure, we divided genes in the human and mouse genomes into three groups (**Supplemental Fig. 2**): single genes (genes present in a genome as a single copy), inparalogs (genes that duplicated from the most similar gene[s] in the genome after the human-mouse split) and outparalogs (genes that duplicated from the most similar gene in the genome before the human-mouse split). This nomenclature has been adopted from (Sonnhammer and Koonin 2002). We distinguished gene families based on the number of inparalogs (genes that duplicated within a species) and outparalogs (from an ancestral duplication). Otherwise the gene family was considered to have only a single gene in a given genome.

In our analyses we explored gene family size, defined as the number of inparalogs/outparalogs, as an important predictor of RT density. The minimal gene family size was two and the maximal number was the maximal size of a gene family in the dataset. Gene families having only a single gene were defined to have size of one (**Table 1**). The RT content (see below) was explored for individual gene family size categories from size one up to ten (gene families of higher size than ten were pooled into a ‘>10’ category). Alternatively, in order to increase the number of gene families within a gene family size category and also to decrease the complexity of the data, we pooled gene families for some analyses into three larger categories: single genes (gene families having a single gene), small gene families (gene families having from two to five inparalogs/outparalogs) and large gene families (gene families having six or more inparalogs/outparalogs). The numbers of gene families within individual gene family size categories are summarized in **Table 1**.

### RT content

The RT data table ‘rmsk’ was obtained from the UCSC FTP server for human and mouse separately (December 2013). The TE data represent the output from the RepeatMasker analysis of the human and mouse genomic sequences. We extracted data for the three most common repeat classes from the ‘rmsk’ table: Long Interspersed Nuclear Elements (LINEs), Short Interspersed Nuclear Elements (SINEs) and Long Terminal Repeats (LTRs). The TEs are classified into subfamilies based on the RebBase classification (Wicker et al. 2007; Kapitonov and Jurka 2008). For our analysis we defined those repeat subfamilies that are lineage-specific based on repeat subfamily names, where the subfamilies unique to one of the two genomes were considered to be lineage-specific. We found 40 LINE subfamilies, 416 LTR subfamilies and 28 SINE subfamilies that are specific to the mouse lineage, and 47 LINE, 266 LTR and 41 SINE subfamilies that are specific to the human lineage (**Supplemental Table 1**).

First, we analyzed RT content (density, abundance and length) with respect to gene family size, as assessed in windows of multiple sizes (10 Kb, 50 Kb, 100 Kb, 500 Kb, 1 Mb, 5 Mb) on each side of a gene. The actual size of the window represents the size after the removal of coding regions of adjacent genes. The density of RTs was then calculated as the proportion of base pairs covered by RTs of a given class divided by the total number of base pairs within the window. The RT densities were divided by the genome-wide averages and, to normalize the data, we took logarithms of base two per (Nellaker et al. 2012). The abundance represents the number of RTs of a group, given the window size. The average length of RTs is calculated as the number of base pairs covered by repeats of a given RT class divided by the RT abundance. The RT content was always compared to the genome-wide average for a given window size (10 Kb, 50 Kb, 100 Kb, 500 Kb, 1 Mb, 5 Mb). Genome-wide averages were based on RT content assessed in sliding windows across the whole genome. All the operations were carried out using BEDTOOLS (Quinlan and Hall 2010) and BEDOPS (Neph et al. 2012) software.

Second, we assessed the contribution by individual RT subfamilies by calculating Shannon’s diversity index using the ‘*Hs*’ method in the ‘*DiversitySampler*’ package (Lau 2012). In order to make the diversity comparable between gene families of various sizes, we used abundances of elements within subfamilies divided by the number of inparalogs in each gene family. The diversity was assessed within 50 Kb windows pooled for each gene family and the numbers of elements were flattened so that no element is present multiple times due to the window overlap.

### RT content vs. gene family size analysis

To test the hypothesis that there is a relationship between gene family size defined as the number of inparalogs/outparalogs and RT content, we analyzed differences in the RT content between individual size categories using the generalized least squares (‘*gls*’) method in the ‘*nlme’* package in the R-project (Pinheiro et al. 2014); and the generalized additive models (‘*gam*’) method in the ‘*mgcv*’ package (Wood 2011). The first method was used to assess detailed differences in RT content between gene families of size from one up to ten and all families of larger size were pooled into one ‘> 10’ category. Gene family size was treated as a categorical variable. We used the full data to explore RT content between genes within a gene family and the interaction between gene family size and window size. Because variances differ among windows of different size we used their z-scores of relative densities with mean zero (genome-wide average) and standard deviation based on the dataset specific to each window size. We modeled the appropriate correlation structure between genes of the same gene family, following which we added two factors: ‘gene family size’ and ‘window size’ and their mutual interaction. We chose the best model using the backward selection procedure and verified individual steps using a combination of hypothesis testing and Akaike’s information criterion (AIC) approach (Akaike 1974). The predicted values and error estimates were obtained using the ‘*predictSE.gls*’ method of the ‘*AICcmodavg*’ package in the R-project (Mazerolle 2015).

The second method was used to describe the relationship between RT content (density, diversity) and gene family size for larger gene families. The common logarithm of gene family size was treated as a continuous variable. To reduce the complexity of the data, we used the RT content within a 50 Kb window that was averaged in a gene family. All the visualization throughout this study was carried out in the R-project using the ‘*ggplot2*’ package (Wickham 2009). The detailed RT densities around chosen gene families were visualized using the Integrative Genomic Browser (IGV; (Robinson et al. 2011).

### Gene Ontology data

Gene Ontology (GO) data were obtained from the publicly available MySQL database (Ashburner et al. 2000; Gene Ontology 2015) downloaded on 05/26/2015). To ensure that there were a sufficient number of gene families associated with a given GO term, we used a flattened set of GO terms based on only the third hierarchical level for all three GO domains (biological process, molecular function and cellular component). This provides sufficient functional distinction, yet includes enough gene families within an individual GO term. We extrapolated GO terms from individual genes onto the whole gene family, assuming that the genes of the same gene family are likely to share the same or similar function. The gene families were pooled into the three gene family size categories (single genes, small gene families and large gene families; see above) and the average RT content for gene families of the same GO term was explored between the three gene family size categories. For our analysis, we considered only GO terms associated with at least five gene families for a given gene family size category. The list of GO terms used in our analysis appears in **Supplemental Table 7**.

### Encode data analysis

We assessed the hypothesis that some of the RTs around the genes of the gene families are involved in gene regulation. To achieve this, we employed open chromatin data based on DNase I Hypersensitive Sites (DHS) from the Encode project (Consortium 2012; Stamatoyannopoulos et al. 2012). We downloaded all DNase-seq datasets available at the ENCODE database for human and mouse before July 2015 (https://www.encodeproject.org). These included DNase-seq datasets by John Stamatoyannopoulos’s lab (UW), Gregory Crawford’s lab (Duke) and Ross Hardison’s lab (PennState). As these data were produced by multiple research groups, we first used a pooling procedure to merge overlapping DHS regions using BEDTOOLS (Quinlan and Hall 2010). Based on the number of tissues/cell types where the DHS region was identified, we created two datasets: DHS1 representing the full DHS region dataset and DHS2 representing tissue/cell type-specific DHS regions identified in less than or equal to ten tissues/cell types. The overlap between DHS regions and RTs in our study was assessed by producing a custom shell script to randomize the location of DHS regions around genes of gene family regions in order to obtain the average random overlap. The significance of the observed overlap was judged by comparison to a randomized distribution.

### Simulation of a second phase of rapid gene family expansion

We simulated a stochastic process of gene duplication and deletion caused by misaligning the non-allelic LINE elements during recombination. Inserted LINE elements underwent slow decay in homology due to mutation. Qualitative properties of the system were assessed. The full description of the simulation algorithm is found in the **Supplemental Methods**. The simulation proceeds by discrete evolutionary time steps (cycles): the algorithm within a single cycle is illustrated in **Supplemental Fig. 8**. Initially, a gene cluster containing five genes interspersed with LINEs is fixed in the population. This resident genotype either stays or, with the arbitrary probability *ι*, is replaced by a new genotype subject to NAHR and further retrotransposition. The time between the population samples is assumed to be sufficiently long to ensure that such a mutant type reaches fixation (i.e. *~4N*_*e*_ generations in the neutral case). At the beginning of the simulation, all LINEs in a cluster are assumed to be nearly exact copies of the transposition-capable elements found in small numbers in both the human and the mouse genomes, and therefore similar to each other. At the end of each cycle, however, and regardless of whether the genotype replacement has occurred or not, all LINEs diverge at a constant rate due to the accumulation of random mutations.

Provided that the replacement type is destined to fixation, the probability of any two LINEs serving as breakpoints for NAHR is calculated as 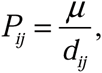, with *μ* being the arbitrary parameter and *d*_*ij*_ being the amount of divergence between the LINEs at the *i-th* and *j-th* intergenic positions, respectively. New LINEs can be inserted at random between the genes, and are assumed to be similar to the LINEs at the beginning of the simulation. The chances of NAHR at particular positions within a cluster increase because the new LINE is assumed to be more similar to any of the old elements than they are to each other. This is reflected by a replacement 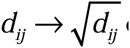 of the amount of divergence at the corresponding positions (see **Supplemental Methods** for details). The simulations were performed in Mathematica 9.0 (Wolfram Research, Champaign, Illinois, USA).

## ACKNOWLEDGEMENTS

VJ and AY were supported from the NextGenProject CZ 1.07/2.3.00/20.0303. VJ was also supported from the Institutional Research Support Grant No. SVV 260 087/2014. CML was supported by the American National Institutes of Health under grants U54 CA143924 and P30 CA23074 and also by Institutional Research Grant number 74-001-34-IRG from the American Cancer Society. We thank the Czech National Grid Infrastructure MetaCentrum running under the program "Projects of Large Research, Development, and Innovations Infrastructures" (CESNET LM2015042) for providing high-throughput computational capacity. We also thank Stephen Proulx for helpful discussion on modelling of the gene family size expansion, Pavel Munclinger in whose lab VJ was based, and Libor Mořkovský for technical support with the UNIX server based at Charles University in Prague.

